# Visual stimuli modulate local field potentials but drive no high-frequency activity in human auditory cortex

**DOI:** 10.1101/2022.07.18.500417

**Authors:** Jyrki Ahveninen, Hsin-Ju Lee, Hsiang-Yu Yu, Cheng-Chia Lee, Chien-Chen Chou, Seppo P. Ahlfors, Wen-Jui Kuo, Iiro P. Jääskeläinen, Fa-Hsuan Lin, Significance Statement

**Author notes:** Corresponding author:* Jyrki Ahveninen, Ph.D., CNY 149, 13th St., Athinoula A. Martinos Center for Biomedical Imaging, Department of Radiology, Massachusetts General Hospital, Charlestown, MA 02129. Authors JA and HJL contributed equally. IPJ and FHL are joint senior authors. Conflict of interest statement*: The authors declare no conflicts of interest.

## Abstract

Neuroimaging studies suggest cross-sensory visual influences in human auditory cortices. Whether these influences reflect active visual processing in human auditory cortices (ACs), which drives neuronal firing and concurrent broadband high-frequency activity (BHFA; >70 Hz), or whether they merely modulate sound processing is still debatable. Here, we presented auditory, visual, and audiovisual stimuli to 16 participants (7 women, 9 men) with stereo-EEG depth electrodes implanted near ACs for presurgical monitoring. Anatomically normalized group analyses were facilitated by inverse modeling of intracranial source currents. Analyses of intracranial event-related potentials (iERP) suggested cross-sensory responses to visual stimuli in ACs, which lagged the earliest auditory responses by several tens of milliseconds. Visual stimuli also modulated the phase of intrinsic low-frequency oscillations and triggered 15–30-Hz event-related desynchronization in ACs. However, BHFA, a putative correlate of neuronal firing, was not significantly increased in ACs after visual stimuli, not even when they coincided with auditory stimuli. Intracranial recordings demonstrate cross-sensory modulations, but no indication of active visual processing in human ACs.

**Significance Statement:** Visual information has a profound influence on auditory processing, particularly in noisy conditions. These “cross-sensory” influences start already in auditory cortices, the brain area that processes sound signals. It has, however, been unclear whether auditory cortex actively processes visual information or whether visual signals only change the way sounds are processed. We studied this question by neurophysiological recordings from 16 participants with epilepsy who had electrodes implanted in their brains due to medical reasons. Using these intracranial recordings, we show that cross-sensory visual information modulates sound processing but triggers no high-frequency activity -- a correlate of local neuronal firing -- in auditory cortex. This result provides important information on the role of sensory areas in multisensory processing in the human brain.

## Introduction

Cross-sensory visual information is known to influence early processing in human auditory cortices (AC) (Molholm et al., 2002; Foxe and Schroeder, 2005; Pekkola et al., 2005; Raij et al., 2010), but the functional significance of these influences remains unclear. According to a more conservative hypothesis, cross-sensory influences play a modulatory role in early ACs, to enhance relevant and suppress irrelevant sound inputs. In line with this suggestion, the effects of unimodal visual stimuli on ACs have often been limited to subthreshold synaptic influences (Schroeder and Foxe, 2002; Ghazanfar et al., 2005; Kayser et al., 2008). For example, laminar extra-cellular recordings in non-human primates (NHP) suggest that cross-sensory visual stimuli modulate the phase of neuronal oscillations but do not increase multiunit firing activity in ACs (Lakatos et al., 2007). At the same time, neurophysiological evidence of *multisensory interactions* (MSI), non-additive changes in neuronal firing rates to multi- vs. unisensory stimuli (Stein and Meredith, 1993), concentrates on higher rather than early sensory areas (Konorski, 1967; Barlow, 1972). One previous NHP model suggested that firing activity is suppressed rather than increased to concurrent audiovisual (AV) vs. auditory inputs in ACs (Kayser et al., 2008).

A more provocative hypothesis suggests that cross-sensory stimuli may be actively processed in early ACs (Calvert et al., 1997; Foxe et al., 2000; Ferraro et al., 2020). This hypothesis has gained support from single-unit recordings in NHPs (Brosch et al., 2005) and other mammals (Wallace et al., 2004; Bizley et al., 2007; Kobayasi et al., 2013), which provide evidence for cross-sensory firing activity in ACs. According to one of these studies, cross-sensory stimuli could drive single-unit firing patterns that convey information of the properties of visual stimuli in the ferret ACs (Bizley et al., 2007). A recent mouse study also suggested firing activity to visual stimuli in the deep infragranular layers of ACs (Morrill and Hasenstaub, 2018).

Whether cross-sensory activity effects such as those seen in animal models occur also in the human ACs difficult to examine using conventional non-invasive neuroimaging. Human neuroimaging evidence of responses to (unimodal) cross-sensory stimuli in ACs consists mainly of studies using MEG/EEG (Giard and Peronnet, 1999; Raij et al., 2010) and fMRI (Calvert et al., 1997; Pekkola et al., 2005), which offer limited means for making inferences of neuronal mechanisms. At the same time, the few previous intracranial studies on cross-sensory influences in ACs have been limited to relatively small samples of participants (N=3-8) and individual-level analyses (Mercier et al., 2015; Ferraro et al., 2020), which may limit the generalizability of results.

Here, we tested the hypothesis that human ACs are not only modulated, but also activated by cross-sensory visual stimuli using intracranial stereo-EEG (SEEG) recordings in 16 participants with depth electrodes implanted near ACs for preoperative monitoring. SEEG provides direct measurements of local field potentials (LFP) from the neuronal tissue, to quantify broadband high-frequency activity (BHFA, above 70 Hz). In contrast to gamma band oscillations that are also visible to MEG and EEG, BHFA signals are believed to be a reliable correlates of local spiking activity (Manning et al., 2009; Miller, 2010; Ray and Maunsell, 2011; Parvizi and Kastner, 2018). Moreover, in studies on ACs, the recording contacts of depth electrodes may extend across the depth of the superior temporal plane, which helps reveal auditory vs. cross-sensory responses at a much greater detail than what is possible with non-invasive recordings, or even with subdural electrocorticography (ECoG). A challenge in previous human SEEG studies of cross-sensory modulation of human ACs has been that the clinically determined anatomical implantation plans vary between participants, thereby complicating robust hypothesis-testing at the group level (Besle et al., 2008; Ferraro et al., 2020). Here, to facilitate group analyses, we complemented traditional “electrode-space” analyses with our recently introduced surface-based source modeling technique that estimates the neuronal activity in the anatomically normalized cortical “source space” (Lin et al., 2021).

## Materials and Methods

### Participants

We studied sixteen participants with epilepsy (15–45 years, seven women) with pharmacologically intractable epilepsy who were undergoing clinically indicated intracranial SEEG recordings. All aspects such as implantation and positioning of electrodes, as well as durations of recordings, were based purely on clinical needs. All participants gave written informed consent before participating in this study. The study was approved by the Institute Review Board Taipei Veteran General Hospital. The details of participants’ demographics and electrode implantation plans are provided in **Table 1**. The sample size was selected to exceed the numbers of participants who were studied in previous comparable intracranial human ECoG (Mercier et al., 2015) and SEEG studies (Ferraro et al., 2020).

**Table 1.** Demographic information for the 16 participants with epilepsy included in analysis. Participants taking part in the additional Experiment 2 have been indicated in the rightmost column. **Abbreviations**: M, male; F, female; OFC, orbitofrontal cortex; MTL, medial temporal lobe.

| Subject | Gender | Age | Seizure onset zones | N of electrodes | Contacts included in analysis | Took part in Experiment 2 |
| --- | --- | --- | --- | --- | --- | --- |
| 1 | M | 17 | Right OFC and MTL | 13 | 83 | X |
| 2 | M | 25 | Left MTL | 12 | 71 | X |
| 3 | F | 29 | Left MTL | 14 | 103 | X |
| 4 | F | 21 | Left MTL | 12 | 70 | X |
| 5 | M | 25 | Right parietal lobe | 11 | 63 |  |
| 6 | F | 45 | Right MTL | 10 | 58 | X |
| 7 | M | 18 | Right MTL | 10 | 68 |  |
| 8 | F | 39 | Bilateral MTL | 10 | 70 | X |
| 9 | F | 16 | Right MTL | 11 | 69 |  |
| 10 | M | 15 | Right parietal lobe | 6 | 38 |  |
| 11 | M | 15 | Right parietal lobe | 16 | 86 |  |
| 12 | M | 38 | Right MTL | 11 | 72 |  |
| 13 | M | 23 | Right MTL | 13 | 94 |  |
| 14 | F | 33 | Right MTL | 14 | 57 | X |
| 15 | F | 23 | Left MTL and insula | 11 | 68 | X |
| 16 | M | 34 | Right occipital lobe | 17 | 98 |  |

### Task and stimuli

In the main experiment, the 16 participants were presented auditory (A), visual (V), or audiovisual (AV) stimuli in a randomized order, at a 1.2–2.8 s jittered inter-stimulus interval (ISI; mean 2 s). In an oddball design, the participant was asked to press a button with their right index finger upon detecting a target stimulus (*p*=0.1) among the non-target stimuli (**Fig. 1**). In the A trials, the non-target stimulus was a 300-ms white-noise burst and the target stimulus an equally long 440 Hz pure tone. In V trials, the non-target stimuli consisted of static checkerboard stimuli (3.5 x 3.5 degrees of visual angle, 300 ms duration) and the target was the same checkerboard but overlapped with a central black diamond pattern. In AV trials, the participant heard and saw the combinations of A and V stimuli; the AV target was the combination of a pure tone and checkerboard with a black diamond. The onset of the V stimulus preceded the A onset by 48 ms. Two six-minute runs of data were collected from each participant. In total, the subject was presented with 100 A, 100 V, and 100 AV stimuli. Auditory stimuli were delivered binaurally via insert earphones (Model S14, Sensimetrics, Gloucester, MA, USA) and the visual stimuli on a 17-inch computer screen (Asus MM17; ASUSTeK Computer Inc, Taipei, Taiwan) positioned at a viewing distance of 60 cm, controlled by using E-Prime (Psychology Software Tools, Sharpsburg, PA, USA).

**Figure 1.**
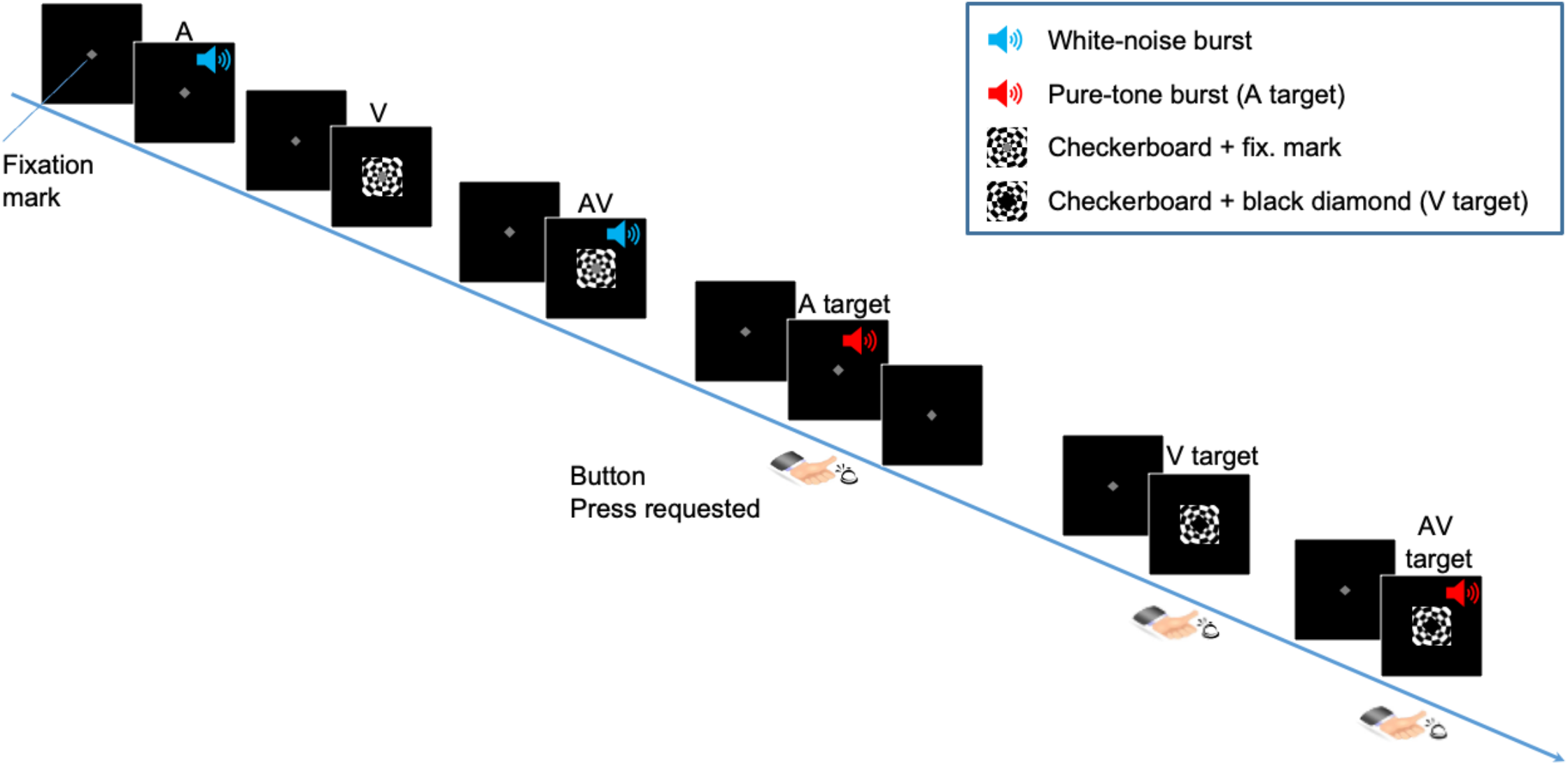
Stimuli and task. Participants were presented with simple A, V, and AV stimuli. In an oddball design, they were asked to press a button upon hearing a rare A, V, or AV target stimulus among frequent non-targets. The non-targets were A noise bursts, V checkerboard patterns, or their AV combinations. The A targets were A pure tones, V targets diamond shapes embedded within a checkerboard, or their AV combinations.

A subgroup of eight subjects participated also in a separate session with more complex task and stimuli than in the main experiment (**Table 1**). This additional session was based on a delayed match-to-sample (DMS) design, with separate lexical and non-lexical parts. The SEEG data from the DMS tasks were analyzed together for the present purposes. In each trial of the <u>Lexical DMS</u>, the participant was first presented an auditory (A) stimulus item, a voice with one of four Mandarin tones. After a randomly jittering delay period of 1 to 1.5 s, the participant was shown a visual (V) probe, which contained one of the possible numbers (i.e., 1, 2, 3, or 4) in the video screen. The participant was asked to press one button with their right-hand index finger if the content of the V probe matched with that of the A item and another button with their right-hand middle finger if it did not match. The task was self-paced, in other words, the task sequence moved on to the following trial only after the participant had responded. In each trial of the Non-lexical DMS, the participant was first presented an A stimulus, consisting of male or female voice stimulus. Then, after a jittering 1-1.5 s delay, the participant saw a V probe, consisting of the Mandarin symbol meaning “male” (i.e., 男) or “female” (i.e., 女). Their task was to press one button with their right-hand index finger if the A voice and V symbol gender matched or another button with their right-hand middle finger if there was a mismatch.

### Data acquisition

SEEG electrodes were placed solely based on the clinical need of each participant with epilepsy, to identify epileptogenic zones. The participants with epilepsy were implanted with from 6 to 17 electrodes (0.86 mm diameter, 5-mm spacing; 6, 8, or 10 contacts per electrode Ad-Tech, Racine, WI, USA). SEEG data were collected at 2,048 samples/s. A white matter contact was used as a reference electrode in five of the 16 participants with epilepsy. In the remaining 11 participants with epilepsy, the scalp EEG electrode at the location FPz served as the initial reference (Caune et al., 2014; Jonas et al., 2014; Rikir et al., 2014; Koessler et al., 2015; Jonas et al., 2016; Mittal et al., 2016; Cam et al., 2017). However, in these 11 participants, the SEEG signals were re-referenced to a contact located in white matter before data analysis. **Figure 2a** depicts the distribution of all electrode contacts of all participants in a standard brain representation.

**Figure 2.**
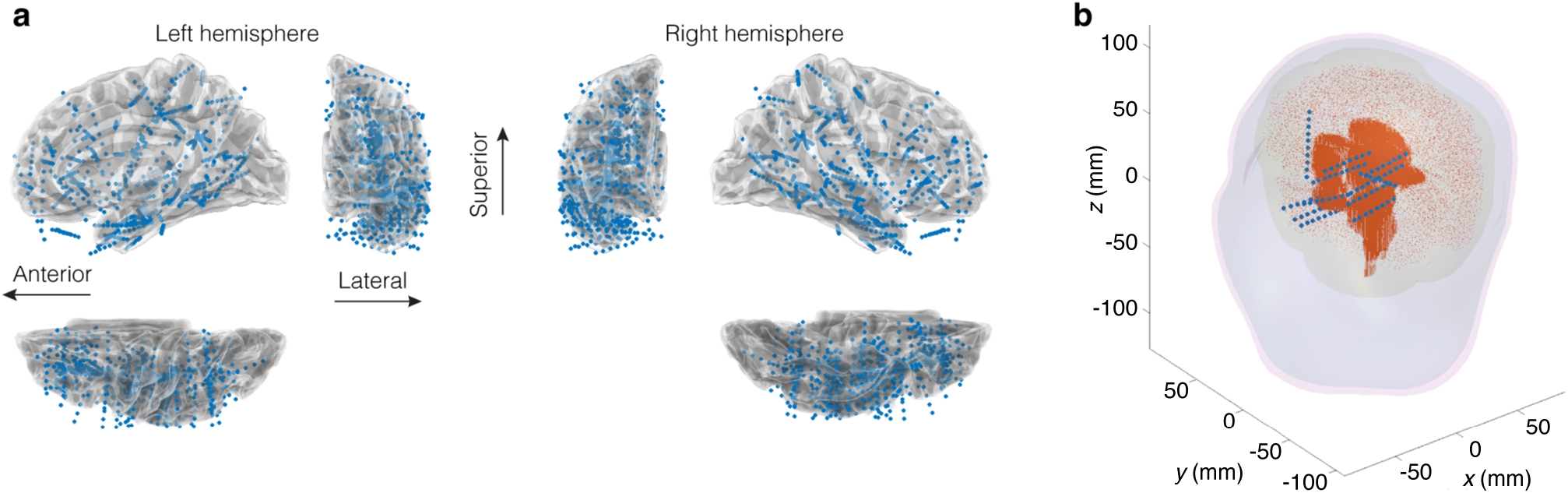
Electrode contact distribution and brain model diagram. (**a**) Locations of all electrode contacts, marked by the blue dots, are depicted in a template brain volume (MNI305). Lateral, frontal, and superior views of each hemisphere are shown. (**b**) A schematic of the source space (orange dots) including cortical and sub-cortical brain areas, locations of the electrode contacts (blue dots), and three anatomical boundaries (inner skull, outer skull, and outer scalp) for the lead field calculation using Boundary Element Model in a representative participant.

Two sets of anatomical MRIs were obtained, one before and another after the implantation surgery of each participant with epilepsy. The pre-operative anatomical MRI was obtained with a 3T MRI (Siemens Skyra, Siemens, Erlangen, Germany) using a MPRAGE sequence with TR/TI/TE/flip = 2530 ms/1100 ms/3.49 ms/7o, partition thickness = 1.33 mm, matrix = 256 x 256, 128 partitions, and FOV = 21 cm x 21 cm. The postoperative anatomical MRI was acquired at 1.5T (GE Signa HDxt system) with a fast spoiled gradient–recalled echo sequence (TR/TE/TI 10.02/4.28/0 msec, flip angle 15°, matrix 256 * 256, bandwidth 31.2 kHz, view 256 × 256 mm, and axial slice thickness 1.0 mm).

### Preprocessing

SEEG preprocessing was performed by using the Matlab (MathWorks, Natick, MA) EEGLAB toolbox (version 14.1.0; http://sccn.ucsd.edu/eeglab/). Electrode contacts that carried excessive line noise were excluded from the analyses based on visual inspection. These analyses also excluded electrode contacts within epileptic lesion sites according to the neurologists (authors H.Y.Y., C.C.C.). SEEG waveforms were notch filtered at 60 Hz to reduce power line artifacts and detrended when cropping the raw data into trials. Trials including epileptiform activity were excluded. SEEG recording was then segmented into epochs of 1500 ms duration with a 500-ms pre-stimulus baseline relative to the A, V, and AV stimulus onsets. SEEG epochs with deflections larger than six standard deviations from the epoch average were discarded. Analyses of time-frequency representations of (TFR) power and inter-trial phase consistency (ITPC) were based on individual trials. The event-related response analyses in **Figs. 3-4** utilize trial-averaged responses.

**Figure 3.**
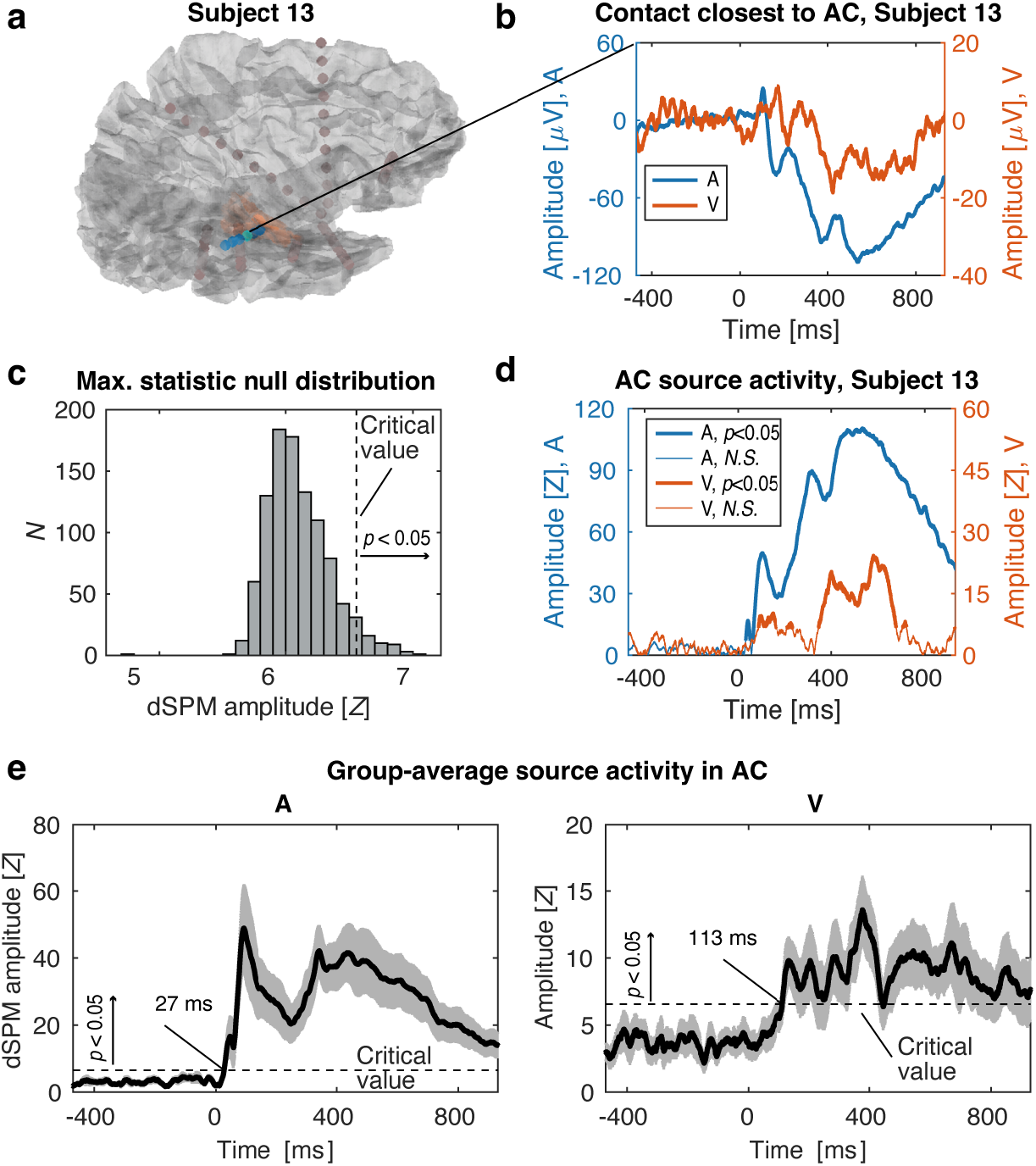
Analysis of trial-averaged intracranial even-related potentials. (**a**) The locations of the superior temporal (sT) electrode contacts (cyan dots) and the location of the right AC ROI are shown in a representative participant (Subject 13) with right-hemispheric implantation. The contact closest to AC appears brighter than the remaining contacts. (**b**) Responses to A and V stimuli in the contact closest to AC in a representative participant. (**c**) Maximum-statistic null distribution, based on 1000 permutations of the group average baseline noise (A, V concatenated) in the AC source estimate. (**d**) AC source estimates in the representative participant (Subject 13). The values above the critical value determined from the group-level null distribution appear on thicker traces for A and V responses (*p*<0.05, maximum-statistic permutation test). (**e**) Group-average AC source estimates to A and V stimuli compared to the critical value determined from the maximum-statistic permutation test. The gray shading refers to the standard-error of mean across the 16 participants.

**Figure 4.**
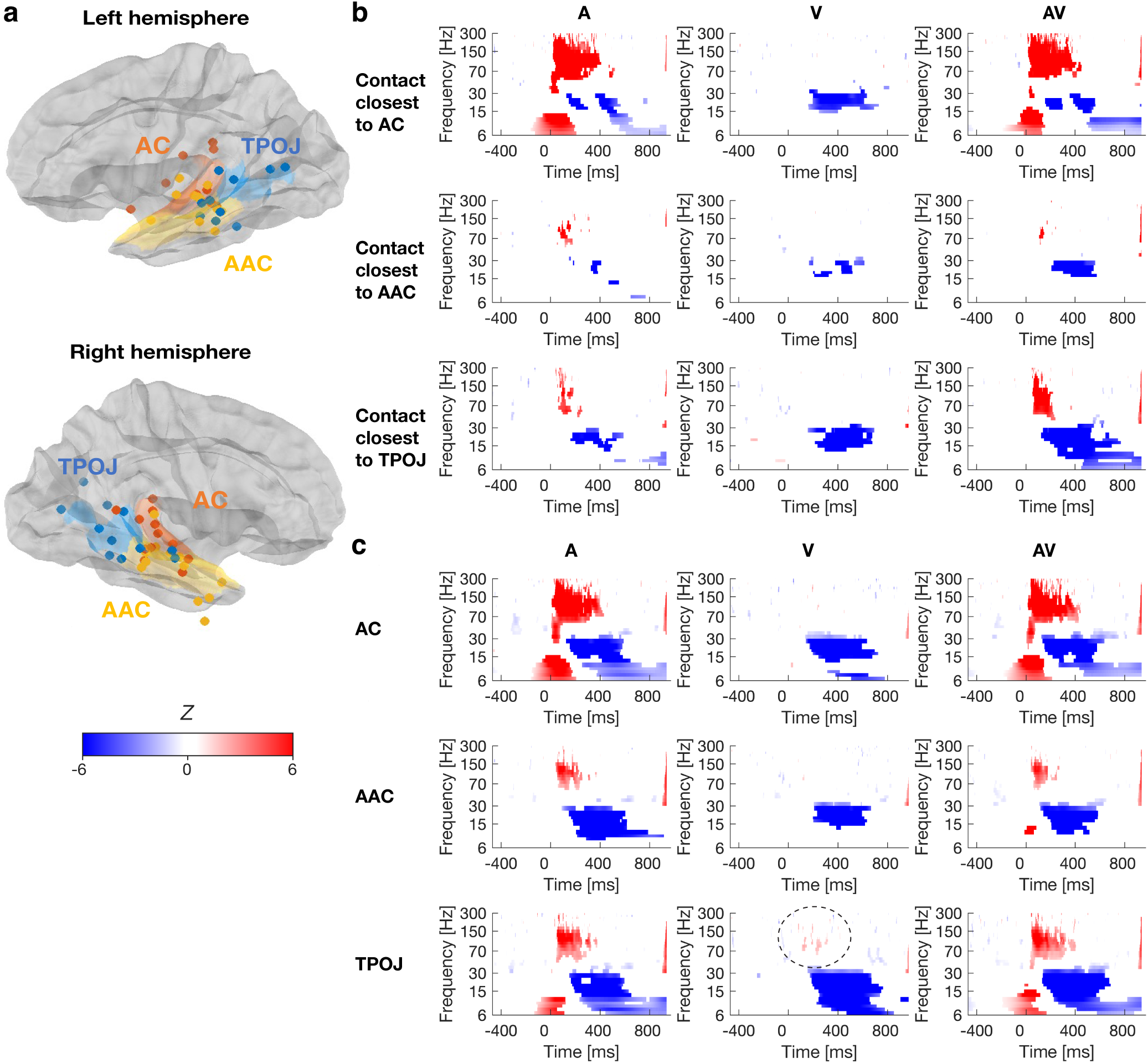
Group analysis of signal power. (a) Analysis sites in the standard brain representation. Each subject’s contacts closest to the HCP-atlas-based ROIs in AC (orange), AAC (yellow), and TPOJ (blue) are marked with orange, yellow, and blue dots, respectively. (b) TFRs to A, V, and AV stimuli in the electrode contacts that were in each subject closest to AC, AAC, and TPOJ. Statistically significant BHFA (>70 Hz, FDR-adjusted p<0.05) emerged to A and AV stimuli only. However, cross-sensory V stimuli suppressed beta (15-30 Hz) power 200 ms after the stimulus onset. (c) ROI analysis of SEEG source estimates. Cross-sensory V triggered suppression of beta and theta/alpha (6-15 Hz) power across the studied ROIs. Traces of weak but significant (FDR-adjusted p<0.05) enhancement of BHFA was observed only in TPOJ (encircled). The TFRs depict the effect size z of the linear mixed effects (LME) model, masked to time/frequency elements with statistically significant effects (FDR- adjusted p<0.05) (Benjamini and Hochberg, 1995). Data from both hemispheres were included a single LME model to determine the statistical significance.

### Source modeling

The coverage provided by depth electrode contacts is sparse and, due to clinical reasons, differs between participants. Thus, it is difficult to analyze the extent of activations within any individual subjects or to conduct anatomically normalized group analyses. These challenges could potentially be addressed by intracranial source modeling of the SEEG data. Although SEEG source modeling has the characteristic limitations associated with ill-posed inverse problems, a crucial advantage compared with MEG/EEG source modeling is the availability of the directly measured depth information of the origins of the signals within the brain parenchyma. Another point is that the BHFA signal that is measurable with SEEG, and which is not visible to non-invasive MEG/EEG, has been reported to be highly focal (Manning et al., 2009; Ray and Maunsell, 2011). Therefore, in addition to conventional signal analyses, we employed inverse modeling of the intracranial source currents to facilitate anatomically normalized group analyses of cross-sensory influences on human AC.

SEEG source modeling was conducted using our recently published strategy (Lin et al., 2021). The locations of electrode contacts in the individual’s brain were identified from the post-surgery MRI, based on discrete dark image voxel clusters caused by the susceptibility artifact at each electrode contact. After specifying the distances between neighboring contacts and the number of contacts in an electrode, the electrode was manually aligned with the dark image voxel clusters in the post-surgery MRI using our in-house Matlab (Mathworks Inc., Natick MA, USA) software with a graphical user interface. Thereafter, contact locations were further optimized (within ±10 mm translation and ±2-degree rotation) by minimizing the sum of squares of image voxel values at all contact locations and their neighboring voxels within a 3x3x3 voxel cube in the post-surgery MRI using the Matlab *patternsearch* function. The SEEG electrode contact locations were then registered to the pre-surgery MRI, which was used to build Boundary Element Models (BEM’s) required for the lead field calculation and to define locations of potential neural current sources. The inner-skull, outer-skull, and outer-scalp surfaces for the BEMs, as well as the cortical source spaces at the gray/white matter boundaries and sub-cortical source spaces, including thalamus, caudate, putamen, hippocampus, amygdala, and brain stem, were automatically reconstructed from the pre-surgery MRI by using FreeSurfer (http://surfer.nmr.mgh.harvard.edu). In each cortical source location, which were separated by about 5 mm from one another, we had three orthogonal neural current dipoles in +*x*, +*y*, and +*z* directions. In sub-cortical regions, sources were separated by 2 mm in the three orthogonal directions. An example of the current source space electrode contacts, as well as of the inner-skull, outer-skull, and outer-scalp surfaces from a representative participant with epilepsy are shown in **Figure 2b**. The lead fields were calculated by using the *OpenMEEG* package (https://openmeeg.github.io/) (Kybic et al., 2005; Gramfort et al., 2010), with the relative conductivity values for air, scalp, brain parenchyma, and skull being 0, 1, 1, and 0.0125, respectively.

The measured sEEG data and the cortical current sources at time *t* are related to each other by **y**(*t*) = **A x**(*t*) + **n**(*t*), where **y**(*t*) is the collection of sEEG data across electrode contacts, **x**(*t*) denotes the unknown current strength, and **n**(*t*) denotes noise. Electrode contacts potentially related to epileptic activity were excluded. The **x**(*t*) has 3 x *m* elements to describe the currents in three orthogonal directions at *m* brain locations and **A** is the lead field matrix. For a unit current dipole source at location **r’** in the +*x*, +*y*, or +*z* direction, the electric potentials measured from all electrode contacts are denoted by **a**(**r’**) = [**a**_x_(**r’**), **a**_y_(**r’**), **a**_z_(**r’**)]. The lead field matrix **A** was obtained by assembling **a**(**r’**) across all possible current source locations: **A** = [**a**(**r’_1_**), **a**(**r’_2_**), …, **a**(**r’***_k_*)], *k* =1, …., *d* where *d* is the total number of current dipole source locations.

To estimate **x**(*t*) using the minimum-norm estimate (MNE), we had ***x***_***MNE***_(*t*) = ***RA**^T^*(***ARA**^T^* + *λ**C***)^-1^***y***(*t*), where ***R*** is the source covariance matrix and ***C*** is the noise covariance matrix ***C*** = 〈***n***(*t*)***n***^***T***^(*t*)〉. The operator 〈·〉 takes the ensemble average across the pre-stimulus interval (-500 ms to 0) with data concatenated across trials. The regularization *λ* tuned the balance between the strength of the estimated neuronal current strength and the discrepancy between the modeled and measured data. We chose *λ* = 10 (Lin et al., 2006).

### Regions of Interest (ROI)

Three ROIs were defined based on the Human Connectome Project (HCP) multimodal parcellation (MMP1) atlas, combined version (Glasser et al., 2016), which was projected to the Freesurfer “fsaverage” standard brain surface representation. From the HCP atlas, we selected the ROIs “early auditory cortex” (AC), “auditory association cortex” (AAC), and “temporo-parieto-occipital junction” (TPOJ), which were subsequently resampled to each participant’s individual cortical surface representation. For the group and individual-level TFR analyses, in each subject, we determined the contact that was closest to the HCP labels AC, AAC, and TPOJ, which were first co-registered to each subject’s cortical surface representation (Arnulfo et al., 2015). For the iERP analysis, average time courses were calculated across the vertices of the ROI, with the waveform signs of sources aligned on the basis of surface-normal orientations to avoid phase cancellations.

### Time-frequency representation (TFR) analysis

TFRs of SEEG power were calculated in Matlab by convolving individual trials with a dictionary of 7-cycle Morlet wavelets. In the case of ROIs of source estimates, the TFRs of power were determined from the sums of squares of the amplitude values along three orthogonal dipole directions. For the group analyses, the power values were then averaged across trials and baseline normalized. In the supplementary individual-level analysis, we compared the power at each time/frequency element after the stimulus onset to the mean power during the respective pre-stimulus baseline period using paired t-tests. ITPC across trials was calculated by dividing the mean amplitudes of the wavelet coefficients across the trials by the mean of absolute values of the amplitudes: 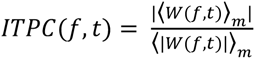 where W denotes the wavelet coefficient, *f* is frequency, *t* is time, and *m* is the trial number.

### Statistical analyses

For the behavioral data, we conducted a repeated-measures ANOVA across the task conditions. We then compared reaction times (RT) and proportions of correct responses (PCR) between AV and A, as well as between AV and V conditions using paired t-tests, corrected for multiple comparisons using the Bonferroni correction (total number of tests = 4). We also calculated behavioral index of multisensory RT benefit, i.e., {(*RT_A_* + *RT_V_*)⁄2} − *RT_AV_* and correlated the result to the AV-(A+V) power contrast estimated from the ROI analysis across the subjects.

For the iERP analysis, a non-parametric maximum-statistic permutation test was utilized to determine a critical value at which the group-average A and V responses significantly exceeded the pre-stimulus baseline. In 1000 permutations, the order of time points of each participant’s baseline period for A and V responses was randomized before calculating the group average. To deal with the multiple comparison problem, from each permutation we selected the maximum amplitude across the A and V baseline periods to be entered to the null distribution. The critical value was determined as the 95^th^ percentile of this null distribution.

TFRs of power values at each frequency were averaged across trials, after which we calculated the base-10 logarithm of each value relative to the average power during the baseline period. Because the ITPC values are bounded within the range between 0 and 1, the values were transformed to Z-score-like values using the Rayleigh test and norminv.m of Matlab (Fisher, 1993). We used a linear mixed effect (LME) model to calculate the statistical inferences of TFRs of power and ITPC in A, V, and AV across participants in both electrode and source analysis using the fitlme.m function of Matlab (Mathworks, Natick, MA). For each condition, the analysis was conducted each element of the relevant time-frequency grid. To compare each task condition separately against the baseline, we used LME models defined by the formula (Wilkinson notation, (Wilkinson and Rogers, 1973)) *M ∼ 1 + (1|subject_id)* where the *M* is the to-be-predicted effect (signal power or ITPC value, respectively), 1 refers to the fixed effect of the intercept, and the term (1|subject_id) refers to the random effect (i.e., random intercept) of the subject identity. Here, the statistical significance was determined based on the fixed effect of intercept. For the contrast between AV and other conditions, we first calculated the to-be-predicted effect size of the contrast of interest, which was then predicted by the same model. The group TFR analysis of Experiment 2 was conducted in the contacts closest to the ROIs representing AC, AAC, and TPOJ, separately for all A item trials and V probe epochs. In this analysis, the A and V epoch data from Lexical and Non-lexical DSM tasks were pooled together. All TFR comparisons were corrected for the inflation of type-I error by controlling the false discovery rate (FDR) across the time-frequency elements.

### Data for reference

Codes used for this study are open to access at https://github.com/fahsuanlin/fhlin_toolbox/wiki. Deidentified data utilized to yield our main results are available at https://dataverse.harvard.edu/privateurl.xhtml?token=d7216368-34bf-4a8a-858a-824b216385fb. Any additional information required to reanalyze the data reported in this paper is available from the Corresponding Author.

## Results

To investigate cross-sensory influences in AC, we presented auditory (A), visual (V), and audiovisual (AV) stimuli to 16 participants with SEEG depth electrodes implanted near ACs for presurgical monitoring. We found strong evidence of subthreshold modulatory effects, reflected as “phase resetting” of ongoing intrinsic low-frequency oscillations and enhanced event-related desynchronizations (ERD) of 15-30 Hz beta oscillations in ACs. Cross-sensory effects were also found in the ROI analysis of iERP source waveforms in the AC. However, although the participants significantly benefited from combining information from the two modalities in terms of the behavioral reaction times (RT), there was little evidence of enhanced BHFA to V stimuli, whether presented alone or in combination with A stimuli in our group analyses.

### Behavioral results

All 16 participants were able to perform the task according to the instruction: On average, they were able to detect almost all targets in auditory (A), visual (V), and audiovisual (AV) conditions, with no statistically significant differences in proportions of correct responses (PCR) between the conditions in our repeated-measures ANOVA (*F*_2,30_=0.45, *p*=0.64). The mean ± standard error (SEM) values of PCR for A, V, and AV target stimuli were PCR_A_ = 0.92 ± 0.04; PCR_V_ = 0.93 ± 0.04 and PCR_AV_ = 0.94 ± 0.03, respectively. The high PCR value for the unimodal V targets provides evidence that the participants were able to focus their gaze according to the instruction. Evidence for significant advantages of combining A and V stimuli were found in RTs to correctly detected target stimuli. The mean ± SEM values of RT for A, V, and AV conditions were RT_A_ = 668 ± 24 ms, RT_V_ = 575 ± 19 ms, and RT_AV_ = 546 ± 24 ms, respectively. There was a significant main effect of task condition (repeated-measures ANOVA, *F*_2,30_=22.7, *p*<0.001). According to subsequent Bonferroni-corrected t-tests across the conditions, the RTs were significantly faster to AV targets than to A targets (*t*_15_ = -5.6, *p*_Corrected_ < 0.001; Cohen’s *d*= -1.5), as well as to AV targets than to V targets (*t*_15_ = -3.0, *p*_Corrected_ < 0.05; *d*=-0.8).

### Intracranial event-related potentials (iERP) in AC

An example of iERP in A and V conditions in the electrode contact closest to AC in a single participant is shown in **Fig. 3b**. The corresponding source estimate for the AC ROI is shown in **Fig. 3d**. Both the closest-contact and the source-modeled waveforms show prominent cross-modal activity in AC in the V condition. Previous non-invasive studies in humans warrant a hypothesis that cross-sensory responses to simple V stimuli are elicited in auditory cortices, lagging the first A responses by several tens of milliseconds (Raij et al., 2010). To evaluate this hypothesis using intracranial SEEG data, we compared the group-averaged source-estimated ROI waveforms calculated to iERPs to A and V stimuli to a critical value determined using a maximum-statistic permutation procedure (**Fig. 3d-f**). The group-average AC source activity ascended significantly above the pre-stimulus noise at 27 ms post stimulus in the case of A responses and at 113 ms in the case of V responses.

### Time-frequency representation of SEEG power

Recent single-unit recordings in non-human primates (Brosch et al., 2005) and other mammals (Wallace et al., 2004; Bizley et al., 2007; Kobayasi et al., 2013), including recent studies in the mouse ACs (Morrill and Hasenstaub, 2018), suggest that instead of modulatory influences only, cross-sensory visual stimuli can also trigger supra-threshold activations in neurons in ACs, reflecting active processing of visual information. Here, we examined BHFA, a putative correlate of multiunit firing activity in the nearby regions (Ray et al., 2008; Parvizi and Kastner, 2018), using both the closest-electrode-contact and source-modeled data.

i. For each subject, we determined the electrode contacts that were closest to the centroids of three cortical Human Connectome Project (HCP) labels across the auditory hierarchy, including “early auditory cortex” (AC), “auditory association cortex” (AAC), and “temporo-parieto-occipital junction” (TPOJ) (e.g., **Fig. 4b**).
ii. A limitation of the electrode-contact analysis is that the recording sites, which were determined by clinical reasons only, differ between the participants. Therefore, to allow for anatomically normalized analyses across the auditory processing hierarchy, we used our recently developed SEEG source modeling approach (Lin et al., 2021). Hypothesis testing was conducted using LME models analogous to those used for electrode-contact analyses in the cortical ROIs AC, AAC, and TPOJ (e.g., **Fig. 4c)**.

TFRs of baseline-normalized signal power values to A, V, and AV stimuli in these contacts of interest and the respective ROIs were then entered to linear mixed effects (LME) models comparing them to the pre-stimulus baseline and determining the differences between AV-(A+V), AV-A, and AV-V contrasts. These LME models assessed the statistical significance of the intercept while controlling for the fixed effect of implanted hemisphere and the random effect of subject identity at all time and frequency instances, with the p-values corrected for multiple comparisons using the false discovery rate (FDR) procedure of (Benjamini and Hochberg, 1995). The critical p value was determined jointly across all conditions/contrasts and contacts-of-interest at 0-500 ms and 6-250 Hz.

#### A, V, and AV stimulus conditions

In the contact closest to the HCP label AC, TFRs to A and AV stimuli showed a characteristic event-related pattern where statistically significant (FDR-adjusted p<0.05) early power increase (or “event-related synchronization”, ERS) at the 6-15 Hz theta/alpha range is followed by a beta-range (15-30 Hz) power suppression (or “event-related desynchronization”, ERD; **Fig. 4b**). At the group level, this theta/alpha-beta ERS/ERD pattern was coupled with a significant (FDR-adjusted p<0.05) sustained increase of BHFA. Interestingly, the beta-range ERD was equally clear to V stimuli alone, suggesting statistically significant (FDR-adjusted p<0.05) cross-sensory modulation of beta-range oscillations in or near human ACs, which started around 200 ms after the stimulus onset. However, no evidence of increases of BHFA, a putative marker of underlying firing activity, were found in the contacts closest to AC, ACC, or TPOJ.

The A, V, and AV stimulus-related TFRs of SEEG power of each individual participant’s contact closest to AC are shown **Fig. 5**. In this individual-level analyses, the single-trial TFR power was compared to pre-stimulus baseline using paired *t*-tests across all trials. The purpose of comparisons against the baseline was also to normalize the TFRs across frequencies to mitigate the 1/f trend of signal power. To avoid false negatives in our interpretation of the lack of BHFA to V stimuli in ACs, we used no correction for multiple comparisons in these specific single-participant analyses. Despite the inclusive criterion, in the electrode-contact-space, evidence of robust increase of visual BHFA, deemed qualitatively different from pre-stimulus noise, was found in only one participant, Subject 8. However, in this participant, the contact that was in the 3D volume closest to early AC appears to be, in fact, located closer to the fundus of STS (a classic multisensory area) than AC, per se.

**Figure 5.**
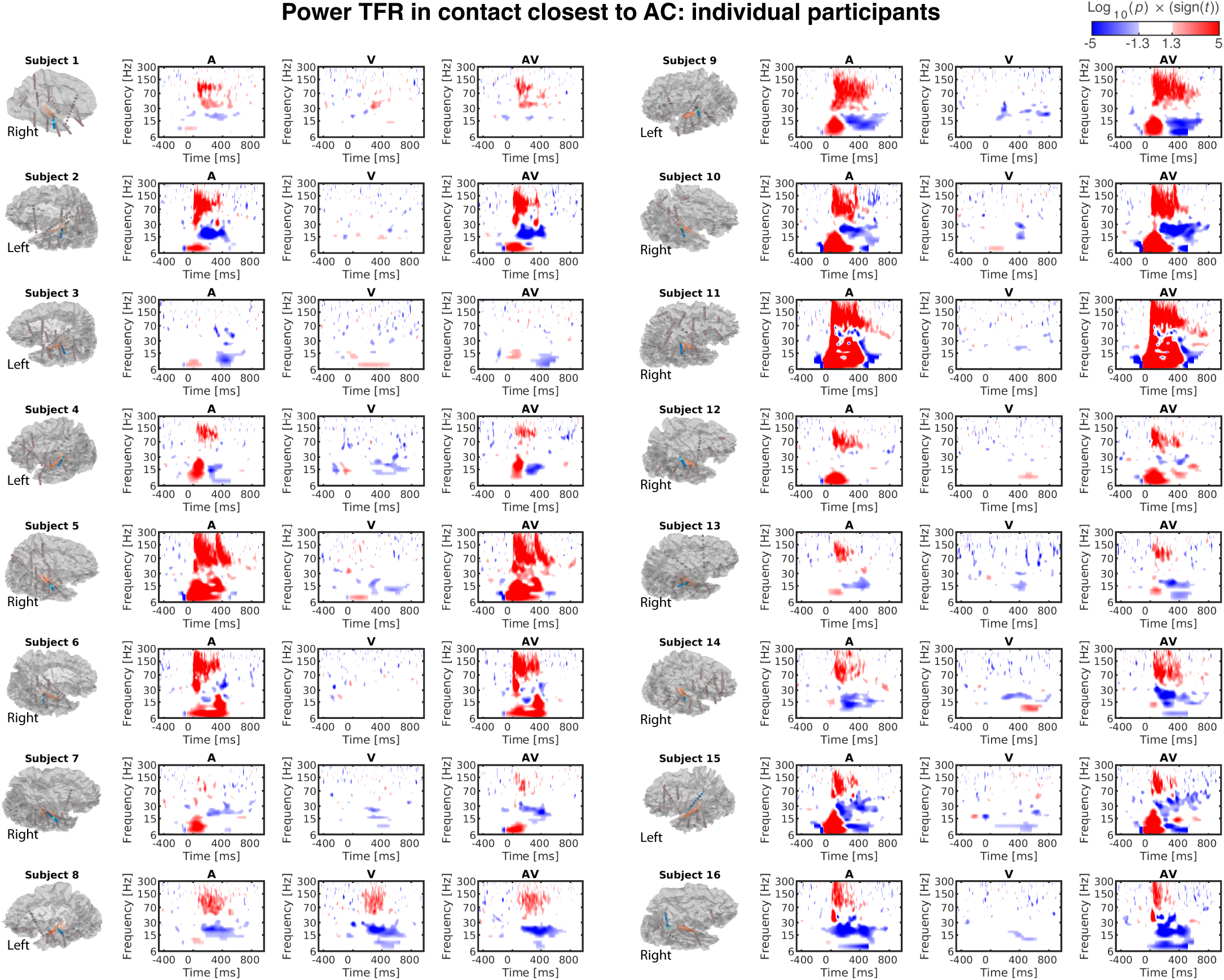
Individual-level TFRs of power. TFRs in the electrode contact closest to AC in individual participants are shown (marked with cyan in each subject; blue dots refer to the electrode where from the contacts was selected). Out of the 16 subject, significant (uncorrected p<0.05) BHF activity was observed in one participant (Subject 8), whose contact closest to AC was in the upper bank of STS, a putative multisensory processing area. The statistical significance of a *t* test relative to the pre-stimulus baseline is shown, thresholded at the uncorrected p<0.05.

Results of our anatomically normalized group ROI source-modeling analysis agreed with the SEEG contact analysis (**Fig. 4c**). At the lower frequency range, responses to A and AV stimuli were characterized by an early theta/alpha ERS (<200 ms, FDR-adjusted p<0.05), followed by a longer-lasting beta ERD (<30 Hz, FDR-adjusted p<0.05). Consistent with the electrode-contact analysis, this later lower-frequency ERD effect was also prominently significant to V stimuli alone (FDR-adjusted p<0.05), constituting the strongest putatively cross-sensory influence across the auditory hierarchy in the present study. As for the BHFA range, statistically significant (FDR-corrected p<0.05) power increases were observed to A and AV stimuli alone in AC, AAC, and TPOJ areas. No significant increases of cross-sensory BHFA were observed to V stimuli in AC or AAC. Traces of significantly increased BHFA to V stimuli were, however, observed in the area TPOJ (FDR-adjusted p<0.05). **Figure 6** shows the source modeling results of the AC ROI in each individual participant.

**Figure 6.**
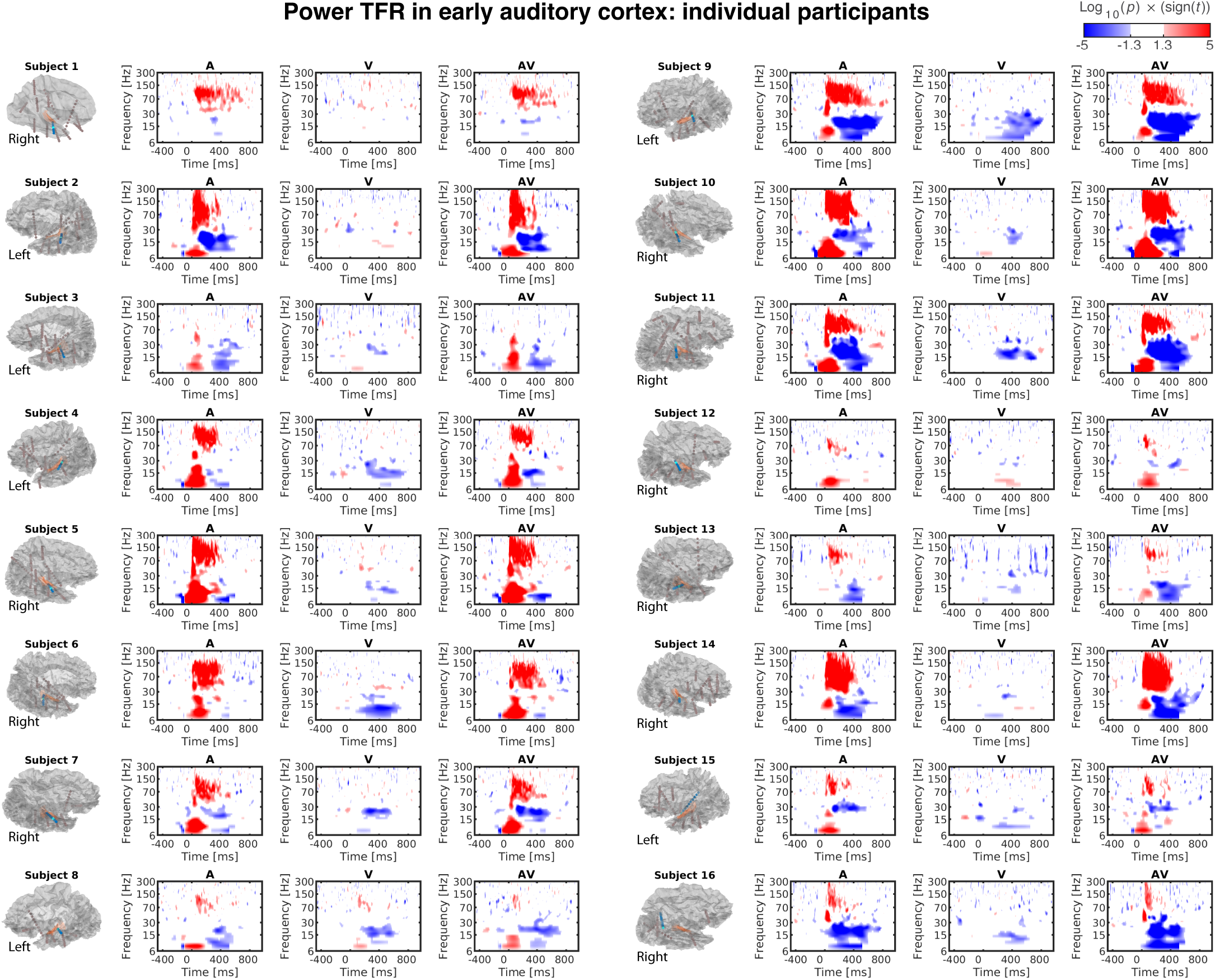
TFRs of power in AC source-modeling ROI in individual participants. The statistical significance of a t test relative to the pre-stimulus baseline is shown, thresholded at the uncorrected p<0.05.

#### Contrasts between A, V, and AV conditions

The group analysis of TFRs from the contact closest to the three ROIs suggested significant (FDR-adjusted p<0.05) non-additive MSI effects, determined from the AV-(A+V) contrast, at the beta range (**Fig. 7a**). As the beta power was suppressed to both A and V stimuli, these positive MSI effects could be viewed as sub-additive reduction of beta suppression. Other than the confirmatory contrast between AV and V conditions, where power changes to a stimulus containing auditory information vs. no auditory information are non-surprising, there was no evidence that the simultaneous presentation of A and V stimuli increases BHFA in the contact closest to AC. Similar to the electrode-contact analysis, the results of source modeling ROI analyses suggested significant (FDR-adjusted p<0.05) non-additive MSI effects in AC, as well as in AAC and TPOJ, which occurred at the theta and beta ranges (**Fig. 7b**). Consistent with the electrode-contact analysis, these positive MSI effects reflect as sub-additive reduction of beta and theta suppression, which occurred to both A and V stimuli alone during the same time/frequency windows. In the AV vs. A contrast, we also observed significant increases of early high alpha activity and subsequent beta ERD effects. Besides the anticipated BHFA effects in the AV-V contrast, the only evidence of increased BHFA was in the AV-A contrast in TPOJ. In particular, no other BHFA effects were observed to AV-(A+V) or AV-A contrast.

**Figure 7.**
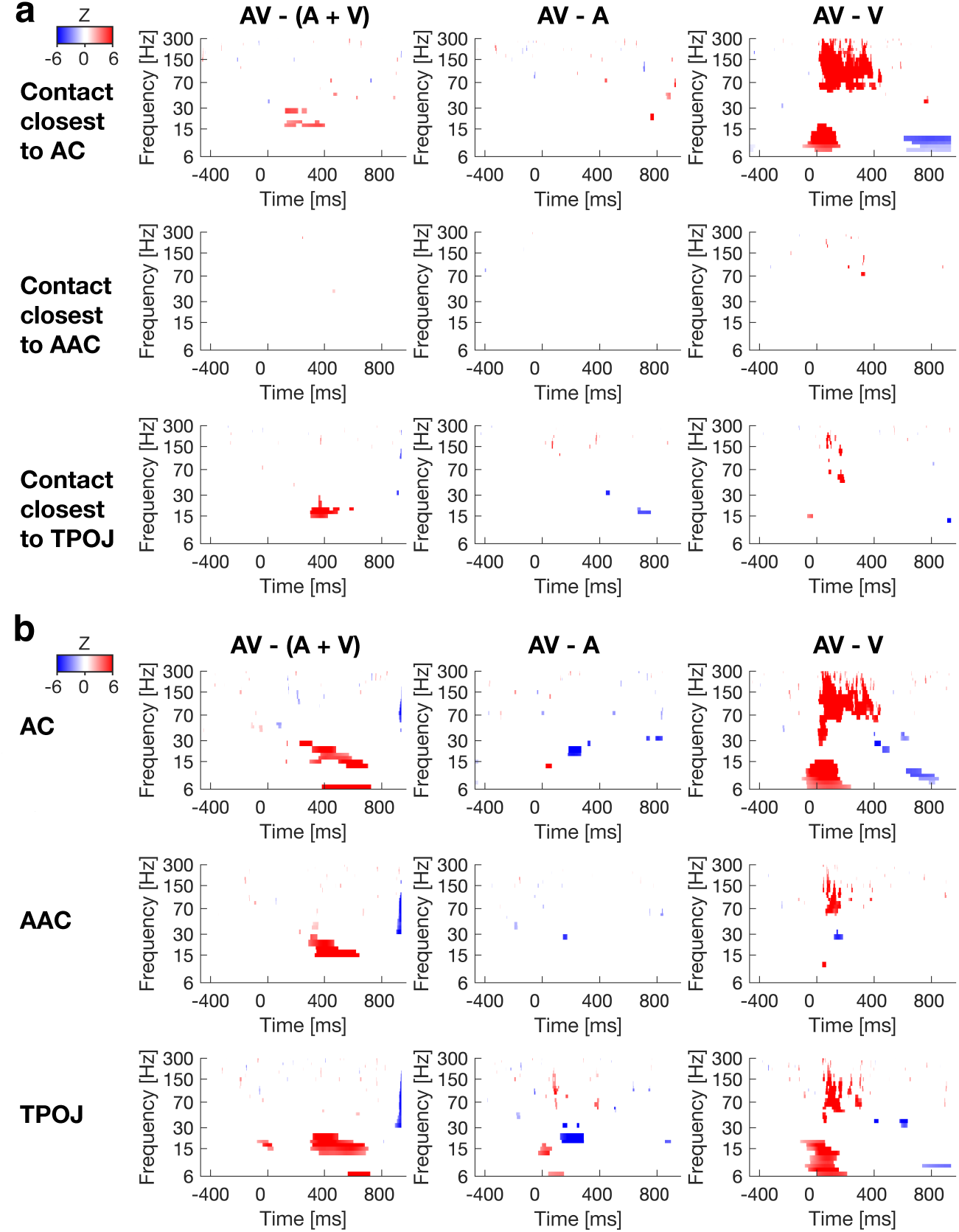
Group analysis of contrasts between conditions using TFRs of power. (**a**) Electrode contacts closest to AS, AAC and TPOJ. Indices of statistically significant non-additive multisensory interactions occur at the beta range in the AV-(A+V) contrast in contacts closest to AC and TPOJ (FDR-adjusted *p*<0.05). The effects in AC occur almost 200 ms earlier than those in contacts near TPOJ. The contrast AV-V shows the anticipated enhancement of BHF and early-latency theta/alpha in AC. (**b**) ROI analysis of source estimates. Indices significant of low-frequency MSIs were observed in all ROIs (FDR-adjusted *p*<0.05). In addition, beta power was significantly suppressed in AC and TPOJ in the AV-A contrast. Apart from the anticipated effects in the AV-V contrast, there was no evidence of effects at the BHFA range. Overall, the effects beyond AC are somewhat stronger than those in the corresponding electrode analysis. The TFRs depict the effect sizes z from the linear mixed effects (LME) model, masked to time/frequency elements with statistically significant effects (FDR-adjusted *p*<0.05) (Benjamini and Hochberg, 1995). Data from both hemispheres were included a single LME model. The ROIs and contacts are shown in the standard brain representation in Fig. 4.

Finally, to evaluate the behavioral relevance of multisensory information, we compared behavioral index of multisensory RT benefit to the AV-(A+V) power contrast estimated from the ROI analysis. No significant correlations (cluster-based randomization test) between the multisensory RT benefit and the AV-(A+V) contrast were observed.

### Intracranial inter-trial phase consistency

In the light of previous studies in non-human primates (Lakatos et al., 2007), we tested whether cross-sensory stimuli affect the phase of ongoing oscillations in human AC. The results suggest significant increases of low-frequency ITPC after V stimuli, both in the electrode contact closest to AC (**Fig. 8a**) and in all three source-modeling ROIs across the auditory processing hierarchy (FDR-adjusted p<0.05) (**Fig. 8b**). In the contact analysis, this ITPC effect was significant already about 100 ms after the onset of V stimuli. Interestingly, the cross-sensory ITPC increases across the auditory processing hierarchy occurred at time/frequency windows where there was no evidence for concurrent increases of TFR power. Apart from the expected difference between AV-A conditions, the ITPC analysis of contrasts yielded no consistently significant differences reflecting potential MSIs in the electrode or ROI analyses.

**Fig. 8.**
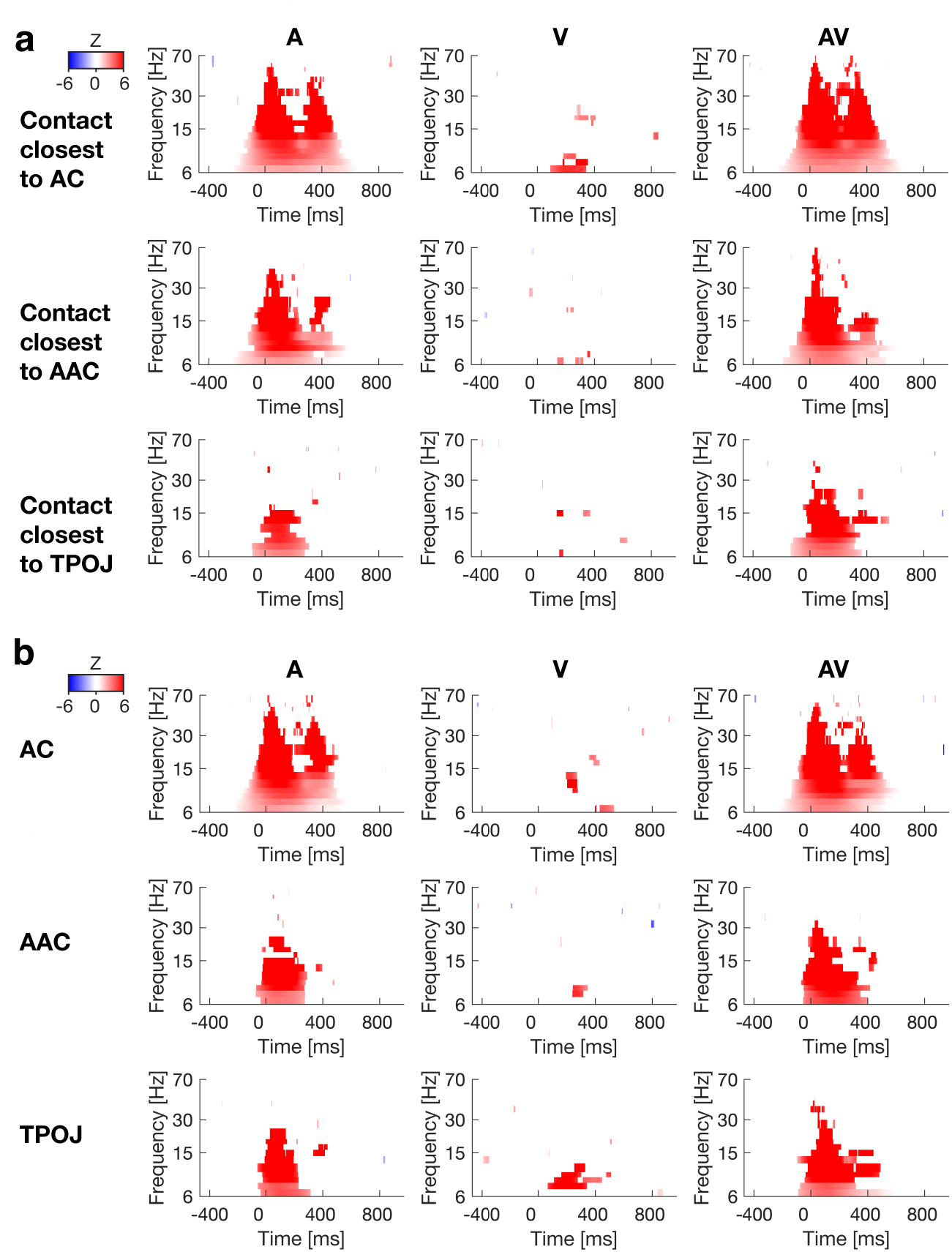
Group analysis of ITPC. (**a**) Normalized ITPC in the electrode contacts closest to AC, AAC, and TPOJ. In addition to the strong ITPC effects to A and AV stimuli, in the contacts closest to AC, there was also evidence of significant increase of low-frequency ITPC to V stimuli (FDR-adjusted p<0.05). These effects started less than 200 ms after the onset of the V stimulus. Most interestingly, the cross-sensory ITPC effect occurred in time/frequency windows where there was no evidence of power increases to V stimuli, in line with the hypothesis that cross-sensory influences could “entrain” intrinsic oscillations of ACs (Lakatos et al., 2007). (**b**) ROI-based group analysis of normalized ITPC values. In addition to responses to A and AV stimuli, there was evidence for significantly increased ITPC to V stimuli across the three ROIs. The data also show the expected early increases of ITPC to A and AV stimuli. The effect sizes z from the LME model have been masked to time/frequency elements where the effects were statistically significant (FDR-adjusted p<0.05). The ROIs and contacts are shown in the standard brain representation in **Fig. 4**.

### Experiment 2: Cross-sensory power changes in auditory cortex during complex stimulation and task

An important concern is whether the results described above, which were obtained during a simple task and basic stimuli, can be generalized to a more complex cognitive setting. Therefore, we analyzed data in a subgroup of 8 participants during additional lexical and non-lexical delayed match-to-sample tasks (DMS, **Fig. 9a**). The behavioral analysis suggested that the participants were able to perform according to the instruction. In the lexical and non-lexical DMS tasks, respectively, the mean ± SEM values of the PCR were 0.92 ± 0.04 and 0.96 ± 0.02. The respective RTs values were 1643 ± 347 ms and 1155 ± 179 ms.

**Figure 9.**
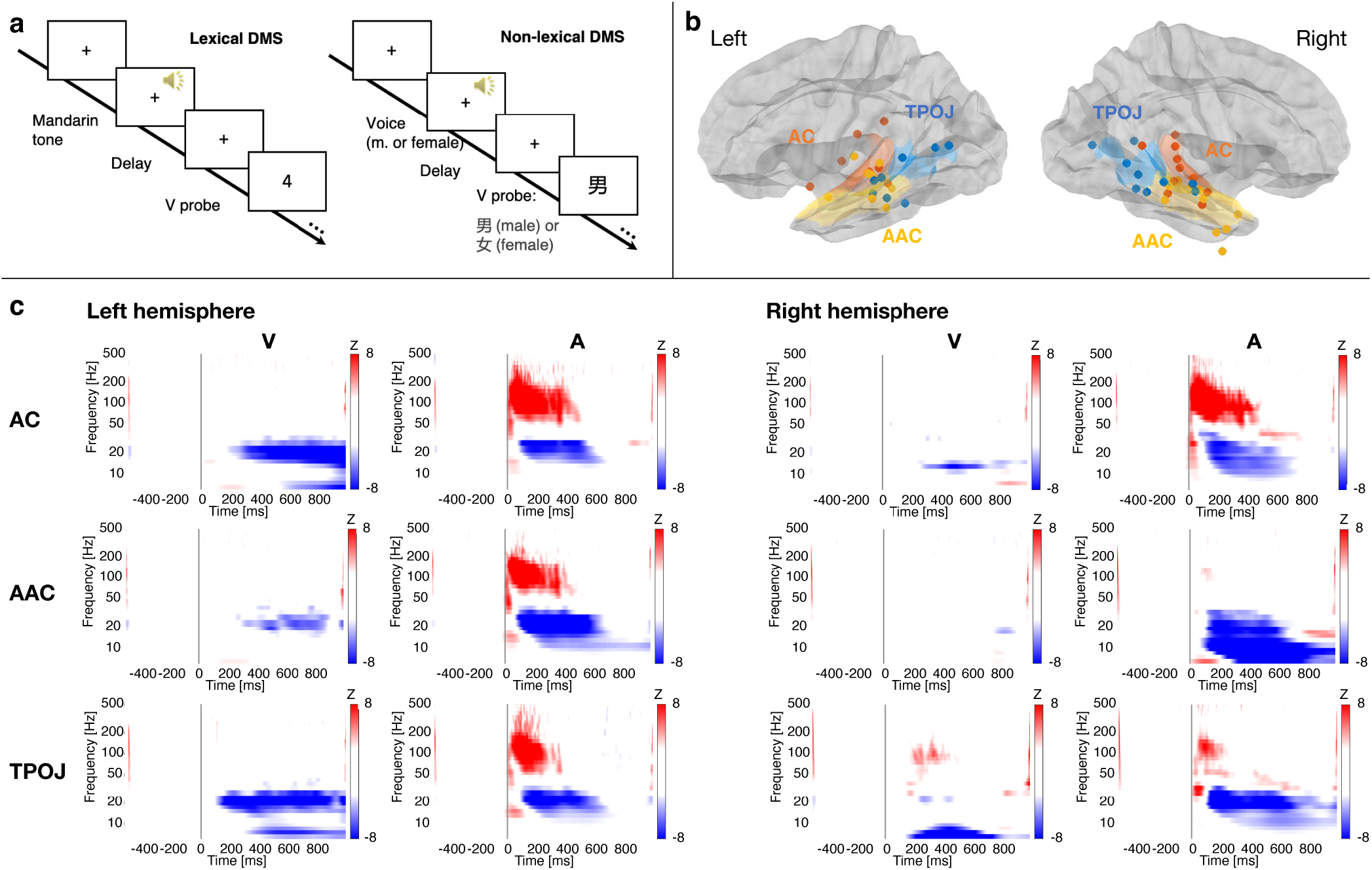
Group analysis of power in contacts closest to AC, AAC, and TPOJ in Experiment 2. (**a**) Delayed match-to-sample (DMS) task paradigms. In each trial of the lexical DMS task, the subjects were presented with an A stimulus, a male or female voice with one of four Mandarin tones, followed by a V probe, a number on the screen. Their task was to identify whether the V probe content matched the tone number of the A. In the non- lexical DMS task, the subjects were presented an A stimulus, a male or female voice, follwed by a V probe, a Mandarin symbol of male or female. Their task was to indicate whether the genders of the A item and V probe matched. (**b**) SEEG group analyses were conducted in contacts closest to AC, AAC, and TPOJ areas. (**c**) Group anaysis of TFRs of power. Statistically significant BHFA (>70 Hz, FDR-adjusted p<0.05) emerged to A stimuli only. In the left hemisphere, cross-sensory symbolic V stimuli suppressed theta (<8 Hz) and beta (15-30 Hz) power, starting roughly 200-300 ms after stimulus onset. Interestingly, these cross-sensory V theta and beta suppression effects were less prominent in the right hemisphere.

**Fig. 9** displays the results of a group analysis, which was conducted across contacts that in each participant were closest to AC, AAC, or TPOJ in both hemispheres. Consistent with the findings in the main experiment, we found no evidence of cross-sensory BHFA increases to symbolic V stimuli that were presented in either lexical or non-lexical contexts in AC, AAC, or TPOJ. At the same time, we observed a significant suppression of theta (<8 Hz) and beta (15-30 Hz) to visual stimuli in contacts closest to the left AC, AAC, and TPOJ. Interestingly, these cross-sensory effects were more obvious in the left than in the right hemisphere. Indices of predominantly left-lateralized effects were observed in the BHFA measures in the higher areas ACC and TPOJ.

## Discussion

Here, we used intracranial SEEG to examine cross-sensory influences in human ACs. In Experiment 1 (*N*=16), we used simple noise-burst sounds and “checkerboard” visual stimuli lacking strong auditory associations, to minimize higher-order (e.g., semantic) feedback influences. The results of Experiment 1 were compared to a cognitively more complex Experiment 2 (*N*=8). Visual stimuli modulated ITPC of low-frequency oscillations in AC, without increasing signal power, suggesting cross-sensory phase resetting of intrinsic oscillations (Lakatos et al., 2007). Visual stimuli resulted in a theta-to-beta-range ERD in ACs. We also observed cross-sensory iERP responses to V stimuli that lagged auditory responses for several tens of milliseconds. However, no robust evidence of visually driven BHFA, a putative correlate of local neuronal firing (Manning et al., 2009; Miller, 2010; Ray and Maunsell, 2011), was found in ACs.

The significant increase in ITPC without concurrent signal amplification, which resembles previous NHP (Lakatos et al., 2007) and human ECoG results (Mercier et al., 2015), could reflect phase modulations of intrinsic oscillations in AC. Lakatos et al. (2007) proposed that such cross-sensory modulations could help ensure that concurrent auditory inputs arrive to ACs at an optimal high-excitability phase, to amplify weak inputs in noisy conditions (Lakatos et al., 2007; Luo et al., 2010; Atilgan et al., 2018). For example, a recent intracranial EEG study suggested that auditory neurons track the phase of visual speech, helping amplify the cortical representation of speech signals (Megevand et al., 2020). Cross-sensory phase resetting of AC oscillations has been reported in non-invasive studies as well (Luo et al., 2010; Biau et al., 2015).

BHFA is thought to reflect a non-oscillatory correlate of neuronal firing (Manning et al., 2009; Miller, 2010; Ray and Maunsell, 2011), offering a measure of supra-threshold activations using human SEEG (Parvizi and Kastner, 2018) (for an alternative model, see (Leszczynski et al., 2020)). In our source analyses, traces of visually triggered BHFA were observed only in TPOJ. In contrast to the strong AV and auditory BHFA patterns, no significant visual BHFA effects were observed in AC, AAC, or in the SEEG contacts closest to these ROIs. In individual participants, significant BHFA to visual stimuli was observed in only one out of 16 participants, whose contact closest to AC was, in fact, located in the upper bank of STS. Consistent with previous smaller-sample human ECoG (Mercier et al., 2015) and SEEG studies (Ferraro et al., 2020), these results suggest that cross-sensory visual stimuli drive very weak, if any, suprathreshold population-level activations near human ACs (see also (Quinn et al., 2014)).

The lack of cross-sensory BHFA in human ACs is in line with neurophysiological recordings in NHPs (Lakatos et al., 2007; Kayser et al., 2008; Kajikawa et al., 2017). Laminar recordings suggested no cross-sensory somatosensory multi-unit activity (MUA) in the NHP primary ACs (Lakatos et al., 2007). Further, while visual stimuli modulated single-unit activity (SUA) and MUA to concurrent sounds, they did not trigger SUA or MUA to visual stimuli alone in the NHP AC (Kayser et al., 2008). Contrasting NHP findings of visually triggered firing effects in ACs have, in turn, been attributed to byproducts of extensive training (Brosch et al., 2005). Beyond this, visually-triggered firing activity has been found in ACs of non-primate mammals only (Bizley et al., 2007; Morrill and Hasenstaub, 2018).

One the most prominent cross-sensory effects in ACs was the visual theta-to-beta ERD, which mimicked the ERD to auditory stimuli. In unimodal studies, alpha-range ERD has been associated with stimulus discrimination/detection (Mazaheri and Picton, 2005): Auditory alpha-ERD increases when a participant comprehends noisy speech sounds (Dimitrijevic et al., 2017) or hears clear rather than degraded speech (Tavabi et al., 2011; Billig et al., 2019). Intracranial sleep studies suggest that auditory alpha/beta ERD occurs only in wakefulness, highlighting its relationship to conscious access to auditory stimulation (Hayat et al., 2022). Consistent with the present results, previous evidence exists that alpha-range ERD is also elicited by crossmodal cues, including visual gestures preceding speech-sound onsets (Biau et al., 2015) and sounds that predict visuospatial target locations (Feng et al., 2017). These cross-sensory ERDs reportedly correlate with behavioral performance in the task-relevant modality (Feng et al., 2017). Theta-to-beta ERDs elicited to visual stimuli in ACs could, thus, reflect crossmodal (or state-dependent (Bimbard et al., 2023)) priming that facilitates auditory-perceptual processing.

Crossmodal visual influences can not only enhance, but also suppress auditory-related fMRI and MEG/EEG signals (Jääskeläinen et al., 2004; Lehmann et al., 2006; Stekelenburg and Vroomen, 2012; Gau et al., 2020). These suppression effects have been associated with predictive coding, enabled by crossmodal information becoming available slightly before the sound onsets (Stekelenburg and Vroomen, 2012; van Laarhoven et al., 2021). An alternative explanation for the visual-triggered theta-to-beta ERD could, thus, be crossmodal suppression of ACs. However, this alternative interpretation is countered by neurophysiological evidence that in the unimodal case, alpha/beta ERD correlates with enhanced rather than suppressed firing activity (Ray et al., 2008).

The pathways for cross-sensory modulations and MSI effects in human ACs have been proposed to include 1) feedback from higher-order areas (Smiley et al., 2007; Luo et al., 2010), 2) direct connectivity between sensory areas (Rockland and Ojima, 2003; Budinger et al., 2006; Bizley et al., 2007; Falchier et al., 2010), and/or 3) subcortico-cortical influences originating in non-specific thalamic nuclei (Hackett et al., 2007). Many of the present effects, including the theta-to-beta ERD and low-frequency MSI effects in ACs, occurred at relatively long latencies. Furthermore, the earliest significant low-frequency MSI effects occurred in TPOJ rather than in AC. Such long-latency effects could reflect supramodal top-down influences (Lakatos et al., 2009) or even behavioral byproducts of the cross-sensory stimuli (Bimbard et al., 2023). However, the cross-sensory ITPC effects in ACs, which became significant already ∼ 100 ms after the stimulus onset, could be early enough to be explained by more direct cross-sensory influences. Notably, stronger early-latency modulations could be expected to visual stimuli with strong auditory associations (e.g., articulatory gestures).

In the classical sense, MSI refers to non-additive changes in SUA, with the AV responses being stronger than the sum of unimodal auditory and visual responses (Stein and Meredith, 1993). Here, we found significant MSI effects only in low-frequency oscillations. However, these MSI effects occurred at time-frequency windows, in which we observed a similar ERD pattern for all three stimulus types: In these cases, the positive AV-(A+V) contrast refers to sub- rather than super-additive influences.

Notably, low-frequency subthreshold modulations (also detectable in non-invasive MEG/EEG) might be easier to detect than cross-sensory driving of neuronal activity. Spiking effect are more focal than modulatory effects in terms of the underlying neuronal tuning (for an AC example, see Eggermont et al., 2011). Furthermore, due to their quadrupolar field pattern, spikes can be recorded near the activated neuron only. The present measure of driving effects, BHFA, could also be harder to detect using sparse SEEG sampling, because higher frequency signals have a shorter “coherence distance” than lower-frequency oscillations. It is also worth noting that BHFA is correlate but not direct measure of underlying neuronal activity (Manning et al., 2009; Miller, 2010; Ray and Maunsell, 2011; Parvizi and Kastner, 2018) (see also (Leszczynski et al., 2020)). Nonetheless, as estimated by the group averages of baseline-normalized power to auditory stimuli, the effect sizes of individual subject’s BHFA were at least as strong as, if not stronger, than the effect sizes of lower-frequency modulatory effects.

The presents analyses are limited to visual cross-sensory influences in or near auditory cortices. This is because of the lack of (clinically determined) SEEG contacts in early visual areas in our participants, a limitation shared by the majority of previous human SEEG studies (Parvizi and Kastner, 2018)(see, however, also (Jonas et al., 2014)). Another limitation relates to MSI effects at the BHFA range. The strongest modulatory cross-sensory effects to visual stimuli in auditory areas was the beta ERD, which started 200 ms after the stimulus onset. A design with a comparable lag between auditory and visual stimuli might have provided more sensitivity for testing this hypothesis.

In conclusion, we found intracranial evidence that cross-sensory visual stimuli modulate sub-threshold LFP signals, including through low-frequency phase resetting in human ACs. However, overall, cross-sensory influences in ACs were somewhat weaker than we had anticipated. As determined via BHFA, there was little evidence of direct activation of ACs by visual stimuli alone. Even when combined with concurrent auditory inputs, visual stimuli drove only weak, if any, increases of BHFA in ACs. Visually triggered BHFA influences were limited to the more polymodal area TPOJ. Finally, many of the most prominent cross-sensory modulatory influences, including the beta-range ERD and concurrent MSI influences, occurred at relatively long latencies in ACs. It thus seems that visual inputs modulate auditory responses, but that little visual information processing takes place in human ACs. Analogously selective attention, such cross-sensory modulatory influences could help suppress irrelevant sound features (Kauramaki et al., 2010), enhance task-relevant rhythmic inputs (Lakatos et al., 2007; Schroeder et al., 2008), and help guide attentional resources to appropriate points of time to enhance auditory speech perception (Zion Golumbic et al., 2013).

## Acknowledgements

Supported by R01DC017991, R01DC016765, R01DC016915; Acad. Finland 276643, 298131, 308431. Russian Science Foundation 22-48-08002.

